# Overactivated epithelial NF-κB disrupts lung development in human and nitrofen CDH

**DOI:** 10.1101/2023.04.07.536002

**Authors:** Florentine Dylong, Jan Riedel, Gaurang M. Amonkar, Nicole Peukert, Paula Lieckfeldt, Katinka Sturm, Benedikt Höxter, Wai Hei Tse, Yuichiro Miyake, Steffi Mayer, Richard Keijzer, Martin Lacher, Xingbin Ai, Jan-Hendrik Gosemann, Richard Wagner

## Abstract

**Background & Objective:** Abnormal lung development is the main cause of morbidity and mortality in neonates with congenital diaphragmatic hernia (CDH), a common birth defect (1:2500) of largely unknown pathobiology. Recent studies discovered that inflammatory processes, and specifically NF-κB associated pathways are enriched in human and experimental CDH. However, the molecular signaling of NF-κB in abnormal CDH lung development and its potential as a therapeutic target requires further investigation.

**Methods & Results:** Using sections and hypoplastic lung explant cultures from the nitrofen rat model of CDH and human fetal CDH lungs, we demonstrate that NF-κB and its downstream transcriptional targets are hyperactive during abnormal lung formation in CDH. NF-κB activity was especially elevated in the airway epithelium of nitrofen and human CDH lungs at different developmental stages. Fetal rat lung explants had impaired pseudoglandular airway branching after exposure to nitrofen, together with increased phosphorylation and transcriptional activity of NF-κB. Dexamethasone, the broad and clinically applicable anti-inflammatory NF-κB antagonist, rescued lung branching and normalized NF-κB signaling in hypoplastic lung explants. Moreover, specific NF-κB inhibition with curcumenol similarly rescued *ex vivo* lung hypoplasia and restored NF-κB signaling. Lastly, we showed that prenatal intraperitoneal dexamethasone administration to pregnant rat dams carrying fetuses with hypoplastic lungs, significantly improves lung branching and normalizes NF-κB *in vivo*. Conclusions: Our results indicate that NF-κB is aberrantly activated in human and nitrofen CDH lungs. Anti-inflammatory treatment with dexamethasone and/ or specific NF-κB inhibition should be investigated further as a therapeutic avenue to target lung hypoplasia in CDH.

## Introduction

Embryonic lung development is a tightly orchestrated process with delicately balanced signaling cues that direct cellular proliferation and differentiation in specific biological phases ^1–3^. These molecular processes are perturbed in patients with hypoplastic lungs due to congenital diaphragmatic hernia (CDH) ^4^. CDH affects 1:2500 newborns of which 30% succumb, while survivors suffer from long term pulmonary and other sequelae ^4,5^. While the diaphragm can be repaired safely ^6^, the high mortality results from lung underdevelopment that features reduced airway branching, alveolar simplification, pulmonary mesenchymal thickening and vascular remodeling ^7^. Nevertheless, the pathobiology of disease related molecular processes in CDH lungs is insufficiently understood, restricting the development of novel therapeutics.

Several animal models to study CDH exist, including the established nitrofen rat model in which oral application of the herbicide nitrofen to pregnant dams at embryonic day (E9) causes CDH in the offspring ^8,9^. Radioactive tracing of orally applied nitrofen to pregnant dams showed that the parent compound reaches the fetal lungs, suggesting that its teratogenicity is mediated through nitrofen itself rather than metabolic intermediates ^10^. The nitrofen model mimics the complete human CDH phenotype with lung hypoplasia (∼100% of pups) and a diaphragmatic defect (∼56% of pups) ^9^. According to a “dual hit hypothesis” that originated from this model, impaired lung branching occurs well before the diaphragmatic defect and external compression to the developing lungs ^11^. Various CDH associated pathways that are involved in fetal lung development have been identified, including retinoic acid signaling or disease specific microRNA or circularRNA profiles ^12–18^. Using unbiased proteomics, we recently discovered that nitrofen-exposed hypoplastic CDH lungs at E21 are enriched in pro-inflammatory processes ^19^. Accordingly, airway basal stem cells derived from human CDH neonates revealed a similar pro-inflammatory transcriptomic and epigenetic signature, with hyperactive nuclear factor-κB (NF-κB; p65). Of note, the inhibition of NF-κB corrected an abnormal CDH associated epithelial differentiation phenotype in human CDH airway basal cells ^20^.

During saccular stages of lung development, aberrant activation of NF-κB in epithelial club cells can later result in impaired alveolarization and maturation in the distal lung ^21^, a key feature in CDH. Moreover, increased NF-κB activation can inhibit canalicular airway branching via interference with Fgf10 ^22^. However, it remains unclear how NF-κB is regulated during abnormal lung development in CDH. In this context, pulmonary macrophages that are enriched in nitrofen CDH lungs could be pro-inflammatory mediators of NF-κB signaling ^23^. Based on these previous observations, we hypothesized that the NF-κB pathway is altered during CDH associated defective lung branching.

Using nitrofen CDH lungs, fetal rat lung explants and human fetal CDH lungs, we show, that the NF-κB pathway is aberrantly increased in CDH lung hypoplasia. Moreover, p65, a transactivation subunit of NF-κB, and downstream inflammatory mediators are enriched in hypoplastic lung explants, as well as in *in vivo* CDH lungs of the nitrofen model. These findings correlate with an increase in NF-κB activation in the airway epithelium of human fetal CDH lung sections and human bronchial epithelial cells after nitrofen exposure. Lastly, we demonstrate that abnormal branching and NF-κB signaling can be rescued with dexamethasone or specific NF-κB inhibition in hypoplastic lung explants *ex vivo* or when applied antenatal to pregnant rat dams *in vivo*.

## Material and Methods

### Study design

Ethical approval was obtained from the health research ethics board of the University of Leipzig (T03-21; TVV34/21) and Massachusetts General Hospital; Boston (IACUC #2022N000003). All animal experiments were performed according to the most current ARRIVE guidelines ^24^. Female Sprague-Dawley rats were housed in the central animal facility of the University Hospital Leipzig and at Massachusetts General Hospital and distributed randomly in individual cages by the animal care staff.

### *Ex vivo* lung explant model

For fetal lung explant studies, pregnant Sprague-Dawley rats were euthanized at E14.5 by pentobarbital-overdose (600mg/kg i.p. ad 1ml 0.9% NaCl). Fetal lungs were collected in HBSS (Thermo Fisher #14170138) and randomly distributed on 0.4µm Greiner Thincerts and placed in a 6-well plate containing 2ml DMEM/F12 (Thermo Fisher #31330038) + 10% FCS (Merck Millipore S0615) + 1% P/S (Thermo Fisher #15140122). Explants were cultured for 2 days with either 4.4mM nitrofen (Cayman Chemicals #20341) dissolved in DMSO or DMSO in medium alone (vehicle) at 37°C in a humidified 5% CO_2_ incubator. We had previously determined that DMSO/medium had no effects on lung explants and therefore is suited as control medium ^19^. Medium was changed after 24h and nitrofen concentration was reduced to 2.2mM for day 2 to account for metabolic processing *in vivo* ^10,11^. After 48 hours, lung explants were harvested and either directly processed for further assays, frozen in liquid nitrogen and stored at -80°C or fixed in 4% PFA and paraffin embedded. For rescue experiments, lung explants were treated with either 1nM dexamethasone (Sigma-Aldrich; #D1756) or 10µM curcumenol (Selleckchem; #S3874) 4 hours after nitrofen exposure on day 0 and again after 20 hours.

### *In vivo* nitrofen CDH model

To induce lung hypoplasia and CDH in the rat fetuses, timed pregnant Sprague-Dawley rats were gavaged with nitrofen dissolved in olive oil (100mg in 1mL olive oil) or olive oil alone (controls) at E9 as described before ^9^. At E14.5 (pseudoglandular stage) and E21 (saccular stage) pregnant rats were euthanized and fetal lungs were harvested. Lungs were randomly selected and either snap frozen in liquid nitrogen and stored at -80°C or formalin fixed overnight and paraffin embedded for further analysis. For *in vivo* rescue with dexamethasone (DXM) pregnant rats were injected intraperitoneally (0.25 mg DXM/kg bodyweight at E10 and E11; and 0.125 mg/kg at E12).

### Human bronchial epithelial cells (BEAS-2B) and human fetal lung sections

BEAS-2B cells were cultured in 50% DMEM (Thermo Fisher #41966052), 50% HamF12 (Sigma Aldrich #N4888) and 1% P/S (Thermo Fisher #15140122) on collagen (Sigma Aldrich, #C5533) coated 6 well plates. 4×10^5^ cells were treated with nitrofen (50µM) for 15min, 1h, 3h and 24h. Treatment with either dexamethasone (1nM) or specific NF-κB inhibitor curcumenol (10µM) was performed for a duration of 24h. Human fetal lung sections from CDH and non-CDH control patients were used from a previously established biobank at the University of Manitoba ^25^. All fetuses succumbed *in utero* or were stillborn at end gestation; non-CDH controls had diseases primarily unrelated to the lung (e.g. placental abruption).

### Western Blot

Tissues and cells were lysed in RIPA Buffer containing protease and phosphatase inhibitors. Protein concentration was determined using BCA Protein Assay Kit (Pierce); equal protein amounts per sample were separated on an acrylamide gel, transferred for 2.5h at 350mA to PVDF-membrane (MerckMillipore) and blocked for 1h in 5% milk in TBS-T (20mM Tris–HCl pH 7.5, 150mM NaCl, 0.1% Tween-20) at room temperature. Primary antibody incubation was performed overnight (s. data supplement for Cat. #) at 4°C. For detection, affinity purified horseradish peroxidase-linked secondary antibodies (Jackson ImmunoResearch) and Western Bright Chemiluminescence Substrate Sirius Detection Kit were used. Densitometry was quantified via ImageJ software.

### RT-qPCR

RNA was isolated with RNeasy Micro Kit (Qiagen); cDNA was generated with MMLV reverse transcriptase (Invitrogen). RT-qPCR was performed using QuantStudio 3 (Thermo Fisher Scientific) with Maxima SYBR Green/ROX qPCR Master Mix (2X). Primer sequences are listed in supplement.

### Immunofluorescence

Paraffin sections (4µm) were dewaxed and dehydrated and washed with TBS. Heat-induced epitope retrieval was performed with Dako Target Retrieval Buffer pH=9,0 for 20 minutes before blocking with 5% goat serum (dianova #053-000-120) or donkey serum (dianova #017-000-121). Immunostaining was completed overnight at 4°C with primary antibodies against p-p65 (S311) and TNF-α (antibodies listed in supplement). Secondary antibodies (Cy3 anti ms/rb) were incubated for 1h at room temperature.

### Imaging and quantification

Microscopy images were obtained with the Zeiss Axio Observer.A1. Image analysis was performed with ZEN and ImageJ. For each staining at least 3 fetal lungs per group from different mothers were used to analyze protein expression. A minimum of 5 images of different regions per lung were taken under identical settings. Corrected total cell fluorescence (CTCF) was calculated as described previously ^26^: Integrated Density - (Area of selected cell x Mean fluorescence of background readings) and divided by area. CTCF per area was summarized as one mean value per sample.

### Statistical Analysis

For statistical analysis, Graph Pad Prism 7 was used. ANOVA analysis, Kruskal-Wallis test, Mann-Whitney-U-Test and Students t-Test were used when appropriate and indicated in the figure legends. Statistical significance was defined as: * = p<0.05; ** = p<0.01; *** = p<0.001; **** = p<0.0001. All graphs are pictured with mean ± SD from at least 3 independent experiments unless indicated otherwise.

## Results

### NF-κB is increased in the developing airway epithelium of human and nitrofen CDH

We had previously discovered that NF-kB is aberrantly activated in cultured airway basal stem cells from human CDH newborns ^20^. To investigate whether these observations in an epithelial cell model from CDH patients reflect the situation in human CDH lungs, we assessed lung sections from CDH (n=3) and non-CDH control (n=4) patients that succumbed before birth (∼40 weeks’ gestation) without any treatment. We found that the transcriptionally active subunit p65 of the NF-κB complex (p-p65) was significantly increased in fetal CDH lungs, particularly in the airway epithelium (Fig. 1A&B). These data demonstrate abnormal activation of NF-κB and important overlap between our cell model ^20^ and *in vivo* lung sections from CDH patients.

**Fig. 1.**
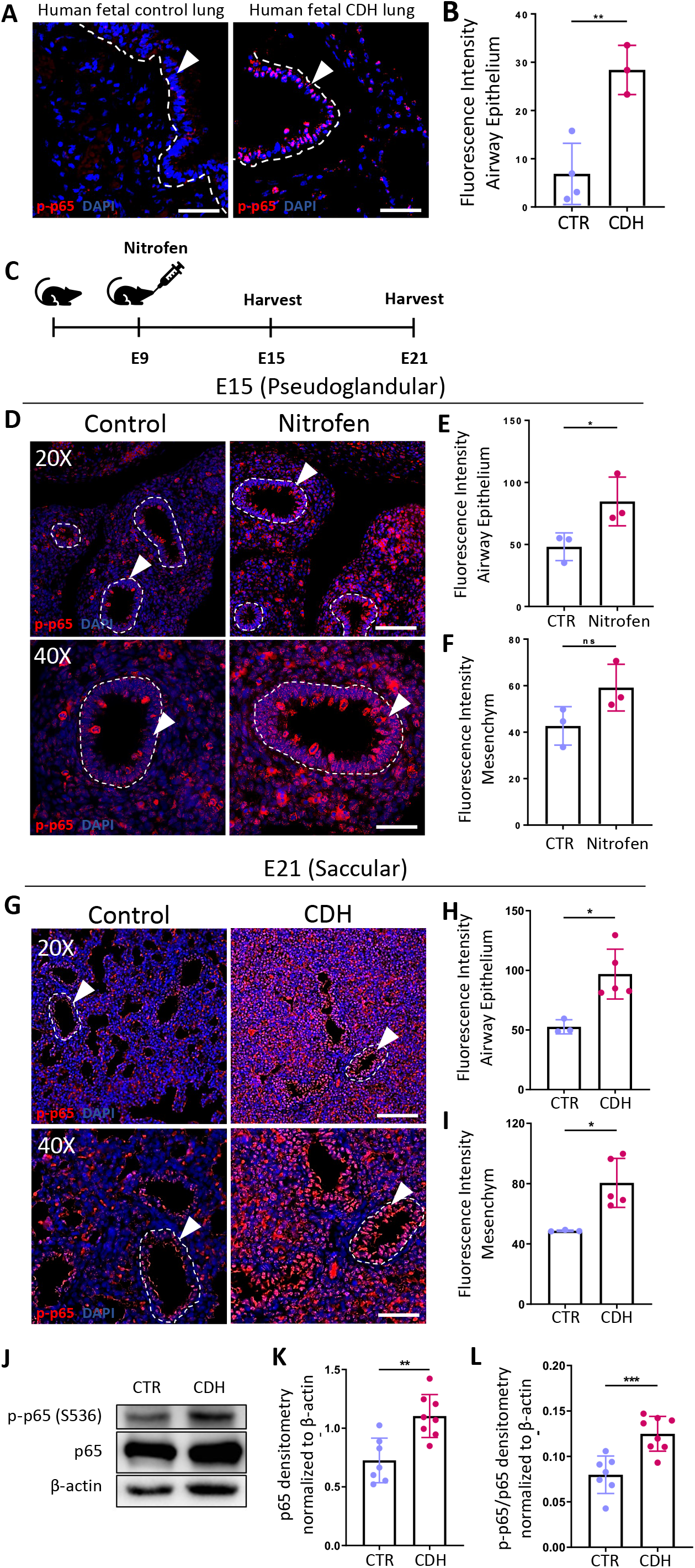
NF-κB is increased in the developing airway epithelium of human and nitrofen CDH. **(A&B)** Representative images and quantification of immunofluorescence staining against phosphorylated p65 (S311) in human age-matched (end gestation) fetal control (n=4) and fetal CDH lungs (n=3). **(C)** Schematic workflow; Nitrofen (100mg in olive oil) was administered per oral gavage to rat dams at E9, (olive oil alone as vehicle controls), lungs were harvested at E15 (pseudoglandular) and E21 (saccular). **(D)** Representative images of immunofluorescence staining against phosphorylated p65 (S311) in E15 control and hypoplastic nitrofen lungs. Airways outlined with dotted lines. Nuclei highlighted via Dapi-staining. **(E&F)** Quantification via corrected total cell fluorescence intensity. **(G)** Representative images of immunofluorescence staining against phosphorylated p65 (S311) in E21 control and nitrofen CDH lungs. Airways outlined with dotted lines. Nuclei highlighted via Dapi-staining. **(H&I)** Quantification via corrected total cell fluorescence intensity. **(J-L)** Representative Western blot and according densitometry for total p65 and phosphorylated p65 (S311) at E21. Airways outlined with dotted lines. Nuclei highlighted with Dapi-staining. For each quantification, at least three 40X images per lung section per group were analyzed and the mean is shown as 1 dot per biological sample. Mann-Whitney U test or unpaired t-test were used when indicated, *p<0,05. 20X Scale bar, 50µm. 40X Scale bar, 100µm. All bar graphs show mean±SD.

To follow up on these findings in human CDH, we explored whether NF-κB signaling is similarly increased in hypoplastic CDH lungs from nitrofen rat fetuses (Fig. 1C). We therefore analyzed lungs of pups at E14.5 (pseudoglandular) and E21 (saccular) via immunostaining and found that p-p65 is increased in hypoplastic + CDH lungs compared to controls at both timepoints, suggesting that a pro-inflammatory signal is maintained in CDH lungs throughout embryonic development (Fig. 1D-I, Fig. S1C-E). Interestingly, p-p65 abundance was especially prominent in the airway epithelium of hypoplastic lungs. In line, mRNA levels of p65 were significantly increased in CDH lungs (Fig. S1B). Western blot analysis confirmed a significant increase in relative protein abundance of both, total p65 and p-p65 in hypoplastic lungs with and without CDH (Fig. 1J-L, Fig. S1F-H), suggesting that this effect is somewhat lung intrinsic and not merely a response to external compression. In line, we detected increased TNF-α in the airway epithelium of CDH lungs at E21 (Fig. S1I&J). Taken together, these findings indicate that the NF-kB pathway and downstream mediators are overactivated and maintained in human and nitrofen CDH.

### NF-κB is aberrantly activated in *ex vivo* nitrofen induced lung hypoplasia

NF-κB is overactivated in human CDH lung epithelium which is reproduced in the nitrofen CDH model. In this model, it is known that nitrofen reaches the fetal lung *in vivo*. To eliminate confounding factors of *in vivo* nitrofen administration (e.g., effects on other maternal or fetal organs, drug delivery efficacy, off target effects) and to mimic CDH associated NF-kB activation *ex vivo*, we cultured fetal lung explants (E14.5 pseudoglandular stage) for 1-2 days with medium alone or medium+nitrofen (Fig. 2A). Thereby, we confirmed that nitrofen exposure results in significantly impaired lung branching with reduction of terminal airway buds and total lung size after 1 and 2 days in culture (Fig. 2, B-D) ^11,19^.

**Fig. 2.**
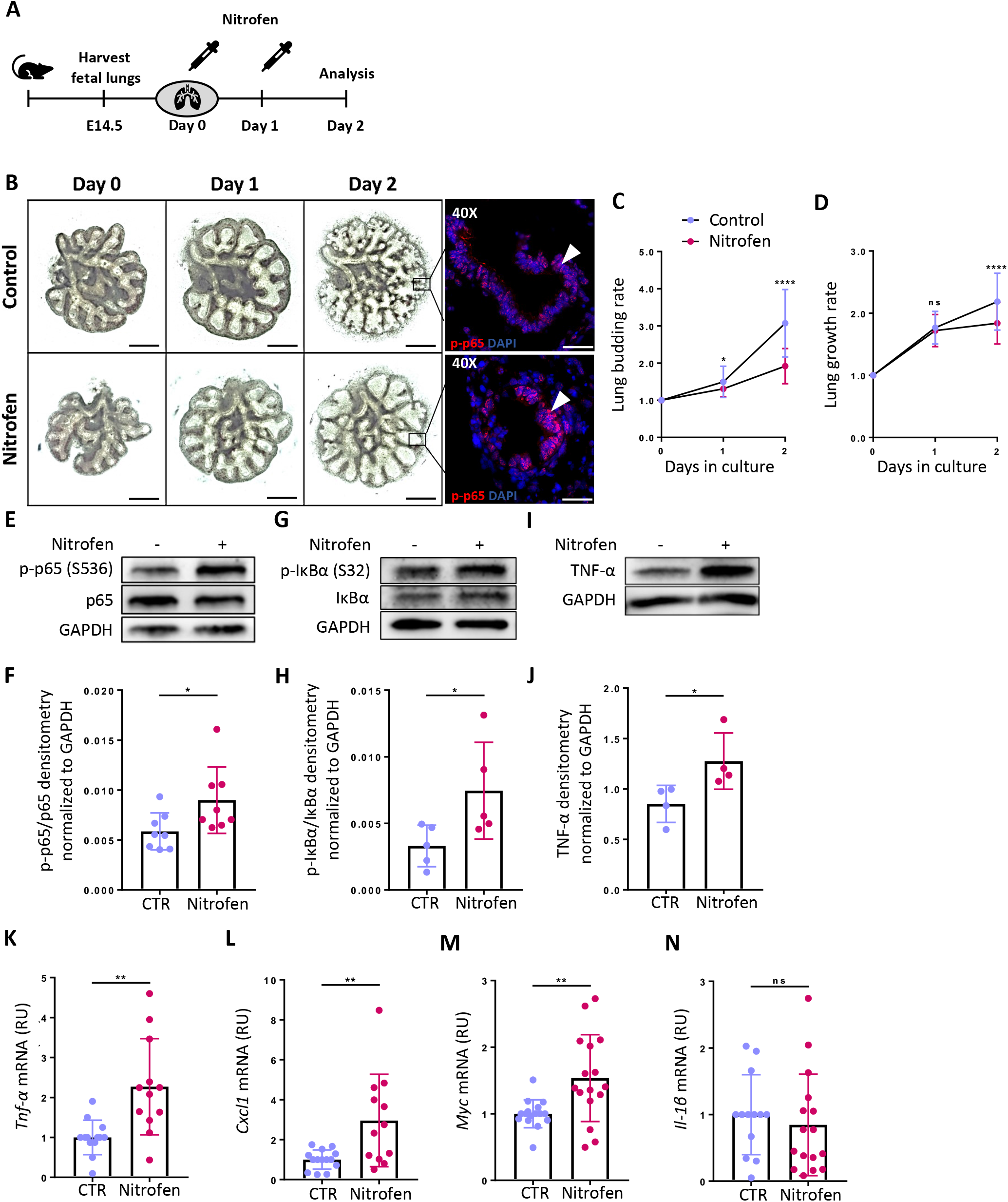
NF-κB is aberrantly activated in ex vivo nitrofen induced lung hypoplasia. **(A)** Schematic workflow; Fetal rat lungs were harvested at E14.5 and cultured for two days in the absence (control) or presence of nitrofen (4.4 mM) to inhibit lung branching. **(B)** Representative images of control and nitrofen exposed lung explants at 0, 1 and 2 days of culture; Immunostaining against p-p65, arrowheads highlight p-p65 expression in distal lung buds. Scale bars, 0.5 mm. **(C)** Quantification of the relative lung budding rate (day2/day0) and **(D)** lung growth rate (day2/day0) of control (n=50-68) and nitrofen (n=52-71) lung explants. Representative Western blot and densitometry analysis in control and nitrofen lungs after 2 days in culture assessing p-p65(S536) (n=8 per group) **(E+F);** phosphorylated IκB (S32) (n=5 per group) **(G+H);** and TNF-α (n=4 per group) **(I+J)**. RT-PCR of NF-κB target genes *Tnf-*α **(K)**, *Cxcl1* **(L)**, *Myc* **(M)** and *Il-1*β **(N)**. Mann-Whitney U test or unpaired t-test were used for statistical analysis when appropriate, *p<0.05; **p<0.01; ****p<0.0001; All bar graphs show mean±SD.

Next, we evaluated NF-κB signaling in hypoplastic lung explants. Immunostaining against p-p65 revealed increased abundance in the distal lung buds of hypoplastic lungs (Fig. 2B). Western blot analysis determined that p65 displays higher phosphorylation in hypoplastic lung explants in culture (Fig. 2E&F). In line, phosphorylation of the inhibitory subunit IκBα and thereby increase in p-p65 activity, and TNF-α protein abundance was similarly increased in hypoplastic lungs (Fig. 2G-J). Downstream p65 transcriptional target genes such as *Tnf-*α, *Cxcl1*, and *Myc* were also significantly increased after 48h of nitrofen exposure when compared to control lungs, while IL-1b was unchanged (Fig. 2K-N). These findings demonstrate that direct nitrofen exposure interferes with lung branching and results in NF-κB hyperactivity during pseudoglandular budding. After we determined that NF-κB is prominently increased in the airway epithelium of human and nitrofen CDH lungs, we asked whether NF-kB is analogously activated after nitrofen exposure in human bronchial epithelial cells (BEAS-2B, Fig. S2A)^16^. Nitrofen in human bronchial cells similarly increased phosphorylation of p65 and IκBα (Fig S2B-D). NF-kB downstream targets were also transcriptionally upregulated in BEAS-2B cells after nitrofen exposure (Fig. S2E-G).

### Dexamethasone can rescue abnormal branching and normalize NF-κB signaling in hypoplastic lung explants and human epithelial cells

After we determined that NF-kB is aberrantly activated in human and nitrofen hypoplastic CDH lungs, we examined whether dexamethasone (DXM), a known antagonist of NF-kB signaling ^27^ that is used to promote lung maturation in premature patients, ^28^ could rescue abnormal lung branching *ex vivo*. We therefore treated *ex vivo* hypoplastic lung explants (E14.5) overnight with 1nM DXM (Fig. 3A). DXM treatment restored lung budding in nitrofen lungs with budding rates similar to control lungs (Fig. 3B&C), and increased mRNA expression of Sonic Hedgehoc associated branching markers such as *Hhip, Gli1* and *Ptch1* which were reduced in nitrofen lungs (Fig. S4). Moreover, DXM treatment decreased p65 phosphorylation in nitrofen lungs back to control levels (Fig. 3D), while total p65 was unchanged, demonstrating inhibitory effects on NF-κB posttranslational modification rather than total protein abundance (Fig. 3D&E). In line, mRNA expression of NF-κB downstream targets *Tnf-*α and *Cxcl1* were normalized after DXM application (Fig. 3F&G). Lastly, we show that DXM treatment can normalize NF-κB activation in nitrofen exposed human epithelial cells (Fig. S3A-F). Thus, DXM treatment restored lung branching in hypoplastic lung explants and normalized NF-κB signaling in nitrofen lungs and human epithelial cells.

**Fig. 3.**
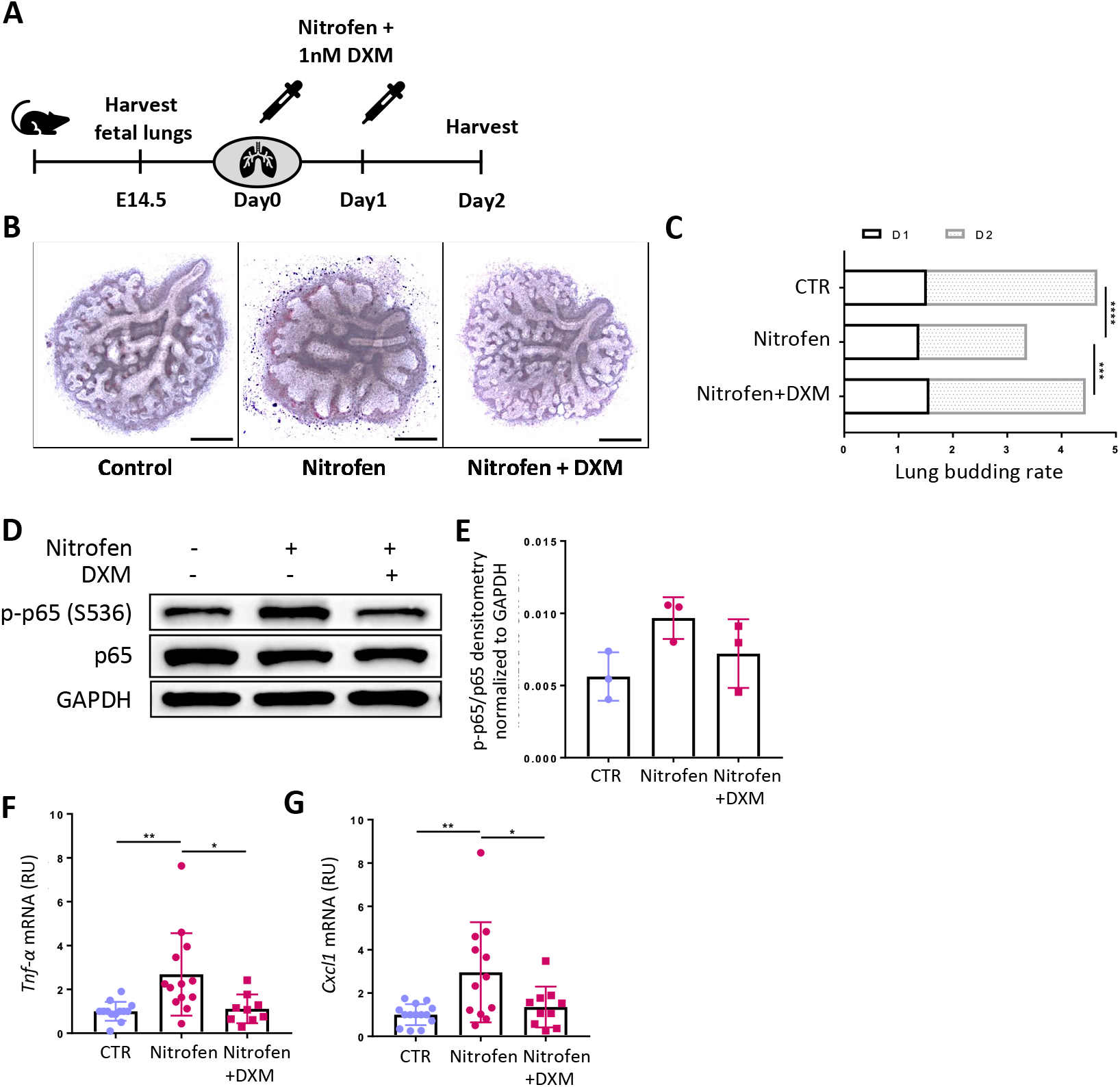
Dexamethasone treatment can rescue nitrofen induced lung hypoplasia *ex vivo* and normalize aberrant NF-κB signaling. **(A)** Schematic workflow; Dexamethasone (DXM; 1nM) was applied 4 hours after initial nitrofen exposure at D0 and again after 20 hours. **(B)** Representative lung explant images from control, nitrofen or nitrofen + dexamethasone treated samples after 2 days in culture. **(C)** Lung budding rates at D1 and D2 from control (n=33), nitrofen (n=34) and nitrofen + dexamethasone (n=40) lungs **(D)** Representative Western blot and **(E)** quantification via densitometry of p65 and phosphorylated p65 (S536) (n=3). **(F)** *Tnf-*α and **(G)** *Cxcl1* mRNA relative gene expression in control (n=13), nitrofen (n=13) and nitrofen + dexamethasone (n=9) groups. Kruskal-Wallis test + Dunn’s multiple comparisons test or ANOVA and Tukey’s multiple comparisons test was used when appropriate, *** p<0,001; **** p<0,0001. All bar graphs show mean±SD.

### Specific NF-kB inhibition can rescue abnormal lung branching and normalize NF-κB signaling in hypoplastic lung explants and human epithelial cells

As glucocorticoids have broad effects aside from NF-κB modulation, we assessed whether specific NF-κB inhibition with curcumenol could restore abnormal lung branching in nitrofen lung explants (Fig. 4). We had previously shown positive effects of a specific NF-κB inhibitor on the abnormal epithelial differentiation of human airway basal cells from CDH patients ^20^. Now, we treated nitrofen and control lung explants with 10µM curcumenol, 4h and 24h after the initial nitrofen exposure (Fig. 4A). This significantly increased lung branching in nitrofen hypoplastic lungs (Fig. 4B&C) and normalized Sonic Hedgehoc associated branching markers *Hhip, Gli1* and *Ptch1* (Fig. S3). While DXM predominantly decreased p65 phosphorylation, specific NF-κB inhibition decreased p-IκBα levels as previously described (Fig. 4D&E) ^29^. Moreover, curcumenol treatment reduced TNF-α mRNA expression and protein abundance (Fig. 4D, F&G). Finally, we show that NF-κB inhibition in nitrofen exposed BEAS-2B cells normalized NF-κB signaling by attenuating IκBα protein phosphorylation and restoring downstream transcriptional targets (Fig. S3G-K). These data suggest, that specific NF-κB inhibition in hypoplastic nitrofen lungs, can rescue abnormal branching and normalize NF-κB activity.

**Fig. 4.**
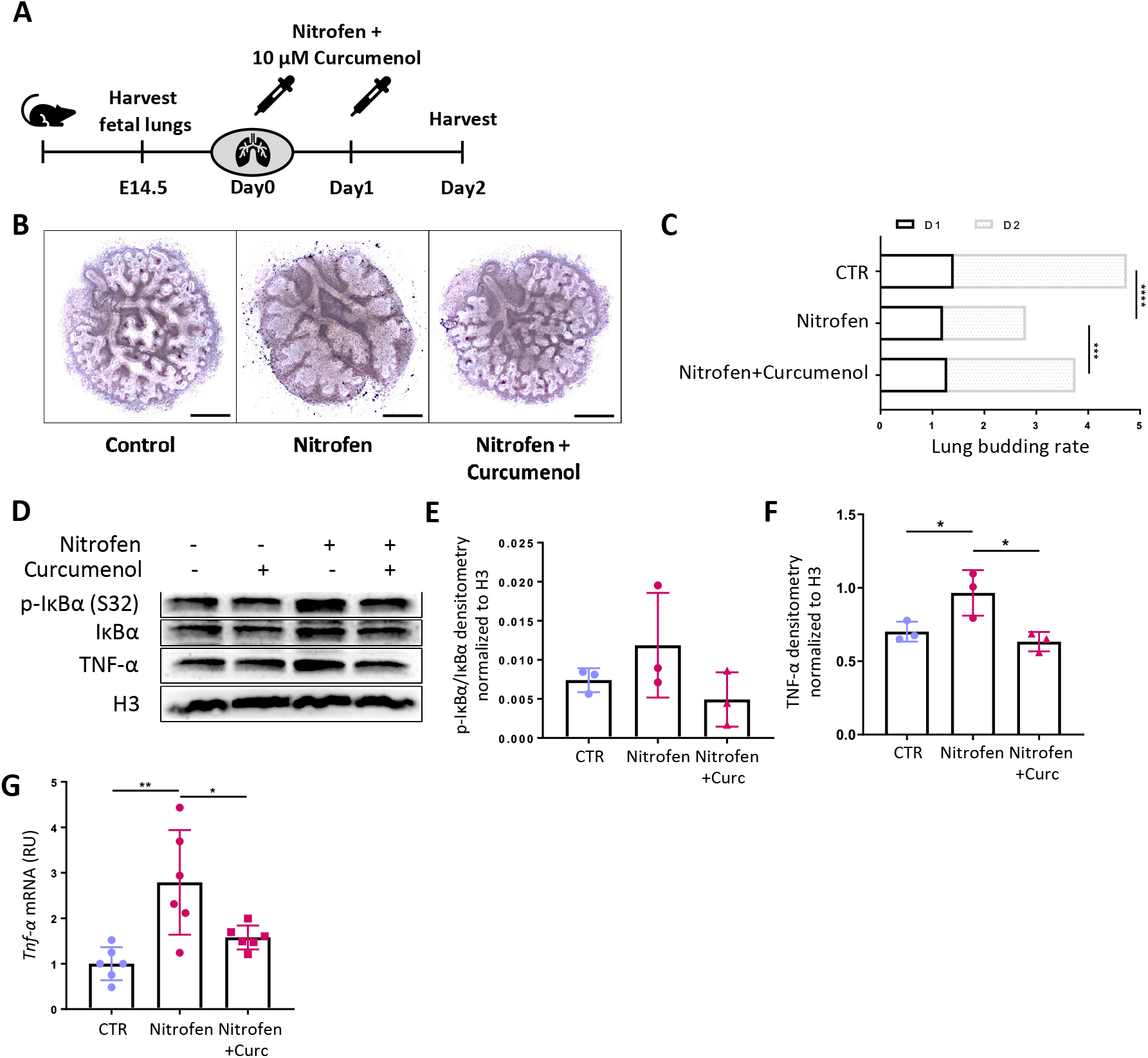
Specific NF-κB inhibition (curcumenol) can rescue nitrofen induced lung hypoplasia *ex vivo* and normalize aberrant NF-κB signaling. **(A)** Schematic workflow: 10µM curcumenol was applied 4 hours after initial nitrofen exposure at D0 and again after 20 hours. **(B)** Representative lung explant images from control, nitrofen or nitrofen + curcumenol treated samples after 2 days in culture. **(C)** Lung budding rates at D1 and D2 from control (n=18), nitrofen (n=17) and nitrofen + curcumenol (n=17) lungs. **(D)** Representative Western blot and **(E&F)** quantification via densitometry of IκB and phosphorylated IκB (S32) and TNF-α (n=3) **(G)** *Tnf-*α mRNA relative gene expression in control (n=6), nitrofen (n=6) and nitrofen + curcumenol (n=6) groups. *Tnf-*α mRNA expression is normalized to actin and control. Kruskal-Wallis test + Dunn’s multiple comparisons test or ANOVA and Tukey’s multiple comparisons test was used when appropriate, *** p<0,001; **** p<0,0001. All bar graphs show mean±SD.

### Antenatal dexamethasone administration *in vivo* rescues impaired lung budding and normalizes NF-κB signaling

Since specific NF-κB inhibitors are not currently used for prenatal therapy, we examined whether early *in vivo* administration of the already prenatally established DXM had beneficial effects on lung hypoplasia in the nitrofen model. Others have previously shown that late gestation DXM treatment improved alveolar development in the nitrofen and sheep models of CDH ^30–33^. Moreover, we showed that DXM can restore abnormal epithelial differentiation and NF-κB signaling in human CDH patient derived airway basal cells ^20^. However, the effects of early (E10-E12) prenatal DXM treatment in nitrofen CDH are unknown. Thus, we administered rat dams that were orally nitrofen gavaged at E9, with intraperitoneal DXM (0.25mg/kg bodyweight at days E10 and E11 and 0.125 mg/kg bodyweight at E12) (Fig. 5A). Nitrofen exposed pups had a significant (∼2-fold) decrease in total peripheral lung buds at E14.5, which is before the physiological closure of the diaphragm. Antenatal *in vivo* DXM treatment markedly restored distal lung budding in nitrofen hypoplastic lungs by ∼ 25% (Fig. 5B-C). and normalized p-p65 abundance (Fig. 5D&E). These data suggest, that the early *in vivo* blockade of aberrant NF-κB signaling in nitrofen lungs improves CDH associated lung hypoplasia.

**Fig. 5.**
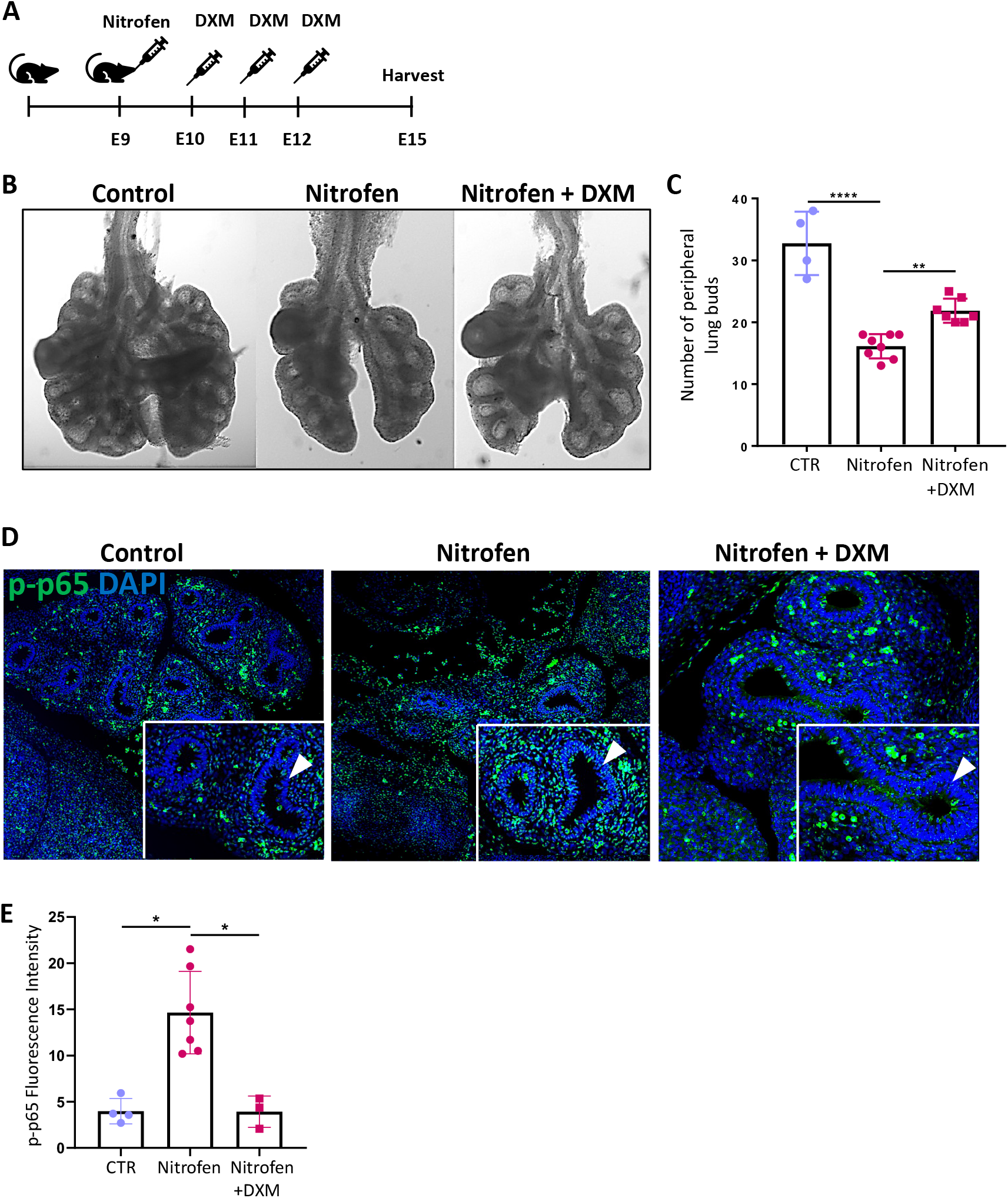
Prenatal dexamethasone treatment *in vivo* increases lung budding in nitrofen induced lung hypoplasia. **(A)** Schematic of the workflow, dexamethasone was applied intraperitoneally at E10, E11 and E12 after *in utero* nitrofen exposure at E9. **(B)** Representative images of control, nitrofen and nitrofen + dexamethasone lungs at E14.5. **(C)** Quantification of number of total peripheral lung buds at E14.5 in control (n=4), nitrofen (n=8) and nitrofen + dexamethasone (n=7) lungs. **(D)** Representative images of control, nitrofen and nitrofen + dexamethasone lungs that were immunostained for p-p65. Nuclei highlighted via Dapi-staining. **(E)** Quantification via corrected total cell fluorescence intensity. 1 way ANOVA and Kruskal Wallis test was used for statistical analysis when appropriate, * p<0,05; ** p<0.01. All bar graphs show mean±SD.

## Discussion

Lung underdevelopment remains the main cause of death in neonates with CDH. While the established nitrofen rat model and other genetic mouse models have revealed different developmentally important pathways that are altered in CDH ^4,7^, more recent reports focused on inflammatory pathways as a common event between animal models and human CDH ^19,20,34^. Building on these previous discoveries we show, in the nitrofen model and human lung specimens from CDH and non-CDH fetuses, that NF-κB is aberrantly activated in CDH lungs. Especially the airway epithelium in nitrofen and human CDH lungs, showed increased p-p65 abundance, which corresponded to enhanced NF-κB pathway activation in cultured human bronchial epithelial cells after nitrofen exposure. Importantly, antenatal blockade of the NF-κB pathway was able to restore abnormal lung branching and normalize NF-κB *in vitro* and *in vivo*.

NF-κB and its downstream pro-inflammatory targets were enhanced in CDH hypoplastic lungs at different developmental timepoints (pseudoglandular and saccular). While NF-κB is ubiquitously expressed in various stages of organ development, it appears that overactivation during early lung development is detrimental, whereas balanced activation postnatally is required for proper alveolar lung maturation ^35^. Accordingly, transgenic overactivation of NF-κB in the airway epithelium of fetal mice at the saccular stage led to distal lung hypoplasia ^21^. Similarly, overactivation of NF-κB ^36^ and associated inflammatory mediators such as IL-1b ^37^, in activated lung macrophages can disrupt airway morphogenesis. In this context, we had previously reported an increase of pulmonary macrophages and monocyte chemoattractant protein 1 in nitrofen CDH lungs, as a potential source of pro-inflammatory signaling in CDH lungs ^23,38^. A more recent and unbiased single nuclear RNA sequencing approach confirmed an influx of macrophages and other immune cells, together with an increase of inflammatory pathways in nitrofen CDH lungs at E21 ^34^. While these studies do not report on the polarity and molecular function of recruited macrophages in CDH lungs, they indicate that immune cells and in particular lung macrophages and their pro-inflammatory effects should be investigated in CDH pathophysiology and potential therapeutic targeting.

While in CDH research, more recent evidence suggests a role of immune cells and pro-inflammatory mediators, this aspect is well characterized in other neonatal lung diseases. In preterm patients, pre- or perinatal inflammation worsens pulmonary outcomes and increases long-term respiratory morbidity ^39,40^. A recent study in preterm primates that were exposed to chorioamnionitis showed that prenatal intrapulmonary inflammation results in significant damage to the fetal lung ^41^. Importantly, the intraamniotic blockade of TNF-α and IL-1b prevents such damage to the developing alveoli ^41^. In line, transgenic mice with increased IL-1b and inflammasome activity in the airway epithelium during saccular lung development had abnormal branching and died shortly after birth ^37^. Other reports in preterm patients determined that pro-inflammatory transcriptomic changes in lung macrophages are associated with increased risk of bronchopulmonary dysplasia ^42^. These results suggest that abnormal activation of inflammatory pathways in the developing airway epithelium can interfere with adequate lung formation.

In addition to our recent findings, emerging evidence supports the observation of enhanced inflammatory signaling in human and experimental CDH. As such, cord blood of human CDH patients sampled directly after birth revealed increased concentrations of pro-inflammatory mediators (e.g.: TNF-α) as the severity of CDH increased ^43^. Here, we identified increased TNF-α abundance in the airway epithelium of nitrofen CDH lungs, which corresponds to a previous report that described increased TNF-α abundance in distal CDH lungs when compared to age-matched non-CDH controls ^44^. In line, we and others previously found increased pro-inflammatory markers (e.g., MCP-1; STAT3) and enrichment of immune cells such as pulmonary macrophages in nitrofen CDH lungs ^19,23,34,45^. Accordingly, different groups have reported positive effects of late gestational anti-inflammatory treatment with dexamethasone (DXM) on distal lung development in the nitrofen rat and a sheep model of CDH ^30–32,46^. Building on these reports, we intended to assess the direct effects of early gestational DXM treatment and specific NF-κB inhibitor treatment on lung branching. Using multiple *ex vivo* and *in vivo* studies, we determined that DXM and specific NF-κB inhibition rescued the budding phenotype in nitrofen lungs, suggesting a functional role of NF-κB during abnormal branching in CDH lung hypoplasia. Despite the convincing results in pre-clinical CDH models, a small prospective trial that analyzed late gestational DXM treatment in human CDH cases did not show beneficial effects ^47^. However, this study was performed in a very small clinical cohort and corticosteroids were administered later during gestation at a relatively low dosage. Promising antenatal surgical interventions, such as fetoscopic endoluminal tracheal occlusion (FETO) ^48^, in combination with established prenatal anti-inflammatory therapeutics such as corticosteroids; or novel experimental approaches (e.g.: extracellular vesicles ^34^) deserve further investigation.

We acknowledge that our study has several limitations. The majority of the data was acquired in the nitrofen rat model. This teratogenic model of CDH, although it mirrors the human condition with all clinical features, is imperfect. We tried to complement our experiments including *ex vivo* lung explants and *in vivo* CDH models, with histological assessment of sections from fetal CDH and non-CDH control lungs, and human bronchial epithelial cell cultures and found significant similarities. However, rigorous studies regarding the efficacy of prenatal targeting of NF-κB are warranted to further investigate its potential benefits, specific molecular effects and potential side effects during lung development. Moreover, our study cannot answer how the pro-inflammatory signal is maintained in hypoplastic CDH lungs well after the *in utero* nitrofen exposure in rats or the teratogenic event in human CDH fetuses.

Our study demonstrates in various models and in human CDH lungs that NF-κB is aberrantly activated at different developmental stages of lung development. Abnormal branching in hypoplastic CDH lungs was rescued by antenatal intervention with dexamethasone or more specific NF-κB inhibitors. Additional experiments that further characterize how these inflammatory patterns are maintained in CDH lungs and what upstream event in human cases prompts this molecular response are underway.

## FIGURE LEGENDS

**Fig. S1. Tnf-**α **is increased in the airway epithelium of nitrofen CDH lungs. (A)** Schematic workflow. **(B)** NF-κB mRNA expression in control (n=7) and nitrofen CDH (n=8) lungs at E21. **(C)** Representative images of immunofluorescence staining against phosphorylated p65 (S311) in E21 control; nitrofen no hernia hypoplastic lungs; and nitrofen CDH lungs. Airways outlined with dotted lines. Nuclei highlighted via Dapi-staining. **(D&E)** Quantification via corrected total cell fluorescence intensity. **(F-H)** Representative Western blot and according densitometry for total p65 and phosphorylated p65 (S536) at E21. *Data points for “control” and “CDH” groups are the same as in Fig. 1. The group “nitrofen no hernia” is for comparison in Fig. S1C-H*. **(I)** Representative images of E21 TNF-α immunostaining in control (n=3) and CDH (n=3) lungs; **(J)** Quantification via corrected total cell fluorescence. Nuclei highlighted via Dapi-staining. For quantification, three 20X images per lung section per group were analyzed and summarized for each biological sample in one dot. Mann-Whitney test was used, *p<0,05. Scale bar, 100µm. All bar graphs show mean±SD.

**Fig. S2. Nitrofen exposure increases NF-κB signaling in human bronchial epithelial cells**.

**(A)** Schematic workflow: BEAS-2B cells were exposed to nitrofen (50µM) and harvested at 15min, 1h, 3h and 24h. **(B)** Representative Western blot and quantification via densitometry for phosphorylated p65 (S536) **(C)** and phosphorylated IκB (S32). **(D)** mRNA relative gene expression in control and nitrofen exposed BEAS-2B cells for **(G)** *Myc* **(H)** *Inf-*β and **(I)** *Cxcl1*. One way ANOVA and Tukey’s multiple comparisons test was used for statistical analysis, *p<0.05; **p<0.01; ***p<0.001. All bar graphs show mean±SD.

**Fig. S3. Treatment with Dexamethasone and Curcumenol can partially normalize aberrant NF-kB signaling in nitrofen exposed human bronchial epithelial cells**.

**(A)** Schematic workflow for dexamethasone (1nM) treatment in control and nitrofen exposed human bronchial epithelial (BEAS-2B) cells. **(B&C)** Representative Western blot and quantification via densitometry for phosphorylated p65 (S536) (n=3). **(D)** *Myc* **(E)** *Inf-*β and **(F)** *Cxcl1* mRNA relative gene expression in control, nitrofen exposed and nitrofen + DXM treated BEAS-2B cells. **(G)** Schematic for curcumenol (10µM) treatment in control and nitrofen exposed human bronchial epithelial (BEAS-2B) cells. **(H&I)** Representative Western blot and quantification via densitometry for phosphorylated IκB (S32) (n=3). **(J)** *Myc* and **(K)** *Inf-*β mRNA relative gene expression in control (n=6), nitrofen exposed (n=3) and nitrofen + curcumenol (n=3) treated BEAS-2B cells. Kruskal-Wallis test + Dunn’s multiple comparisons test or ANOVA and Tukey’s multiple comparisons test was used when appropriate, *** p<0,001; **** p<0,0001. All bar graphs show mean±SD.

**Fig. S4. *Hhip expression* is decreased in hypoplastic lungs and normalized after dexamethasone and specific NF-kB inhibition**.

mRNA relative gene expression in control (n=14), nitrofen (n=14) and nitrofen + dexamethasone (n=8) groups for **(A)** *Hhip*, **(C)** *Gli1*, **(E)** *Ptch1;* mRNA relative gene expression in control (n=14), nitrofen (n=14) and nitrofen + curcumenol (n=5) groups for **(B)** *Hhip*, **(D)** *Gli1*, **(F)** *Ptch1*.

## Notes

Funding: This work was supported by the German Research Foundation to RW (461188606).

### Competing Interest Statement

The authors have declared no competing interest.

