## Supplementary figures and images for "Overactivated epithelial NF-κB disrupts lung development in human and nitrofen CDH"

### Supplemental Material

#### Supplemental Figure 1

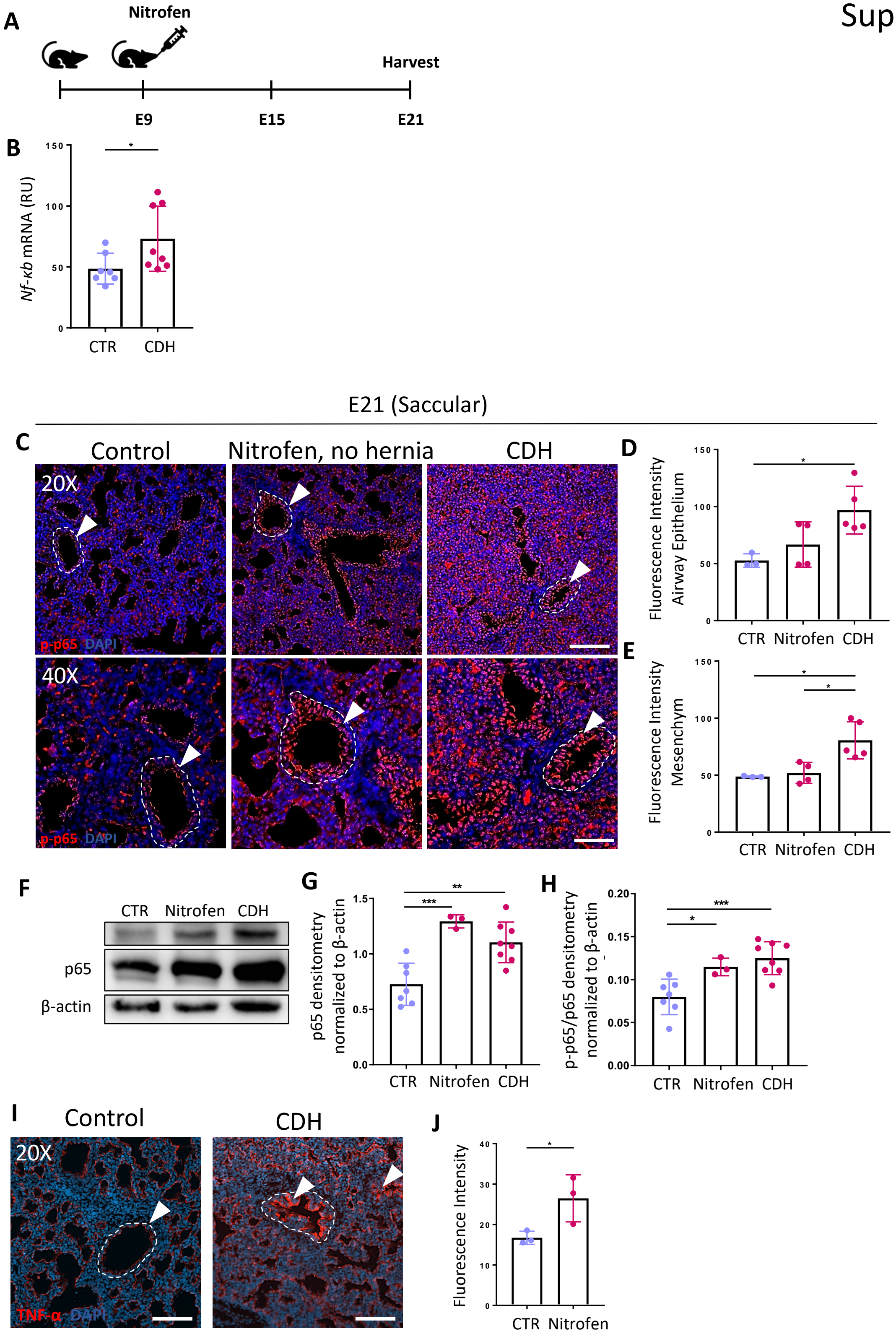

#### Supplemental Figure 2

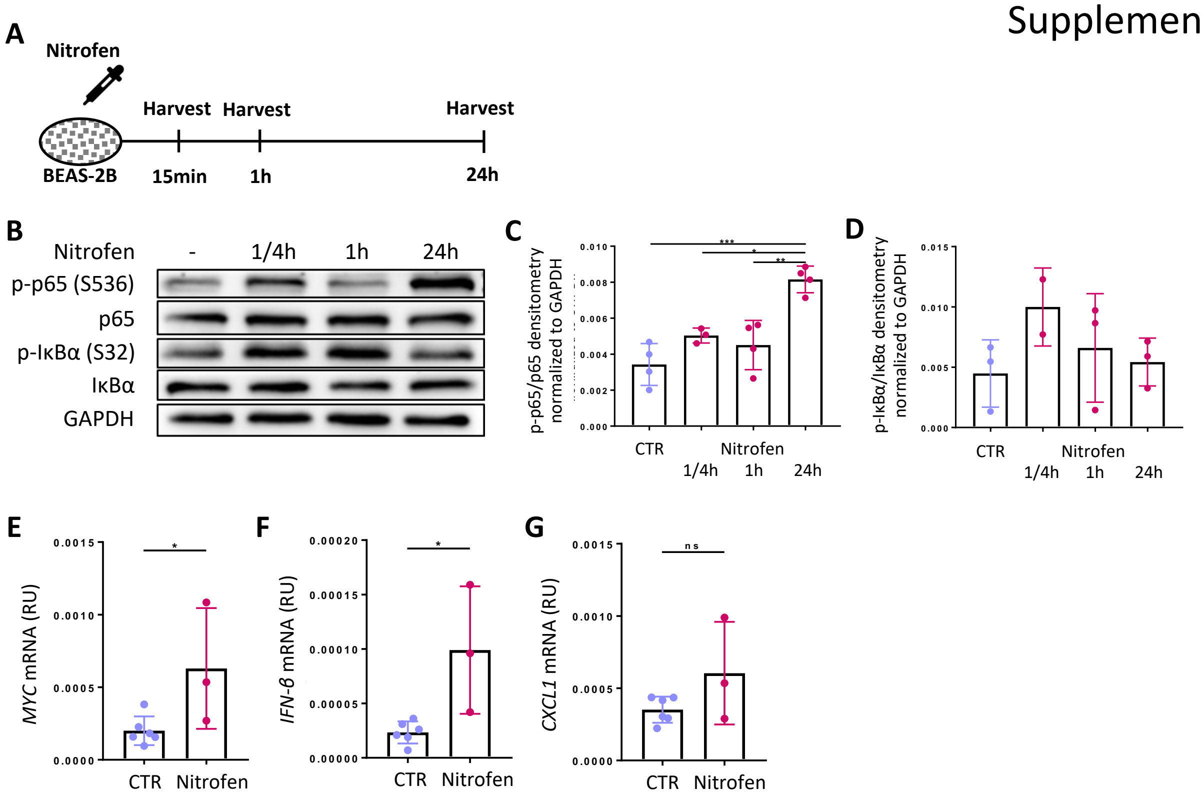

#### Supplemental Figure 3

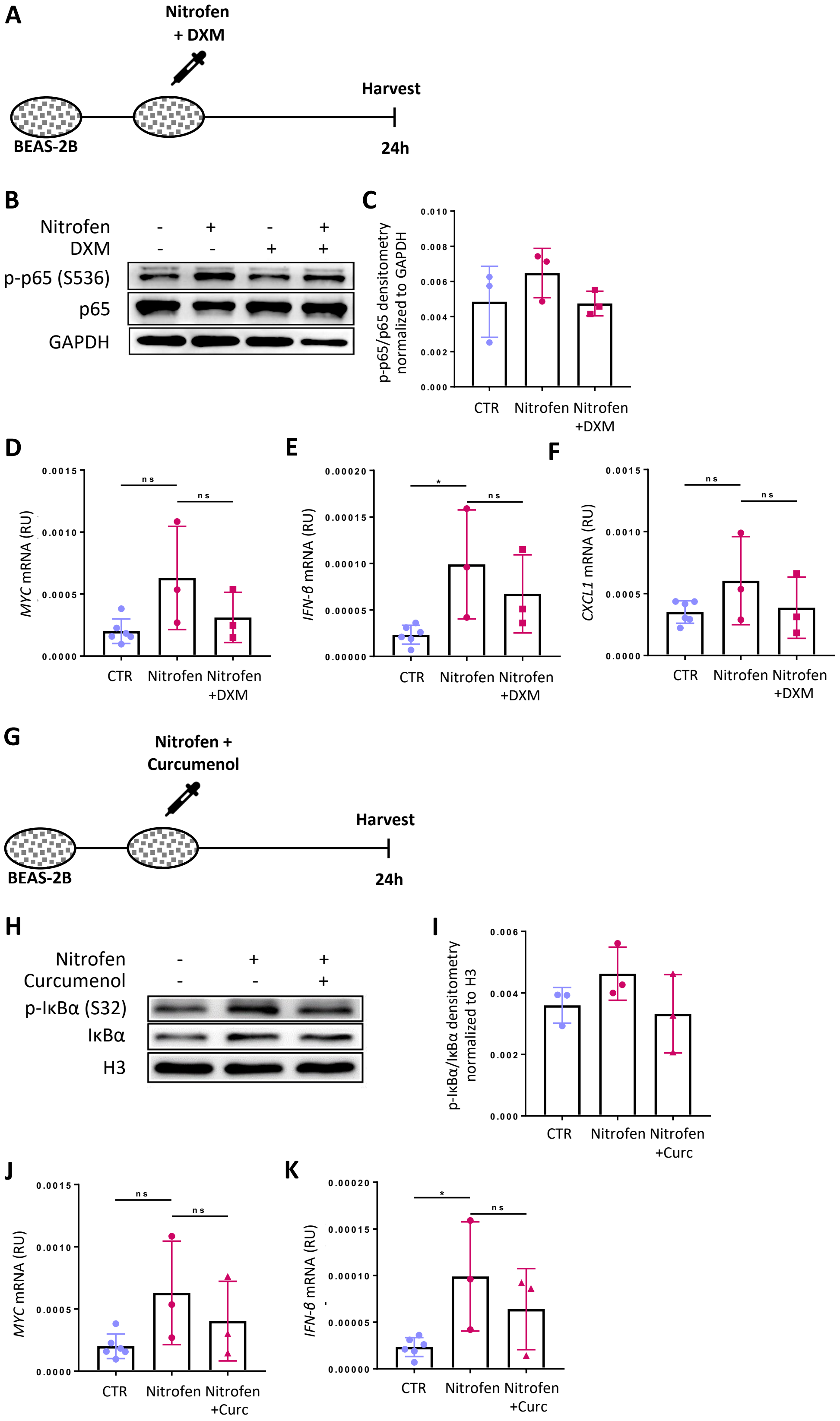

#### Supplemental Figure 4

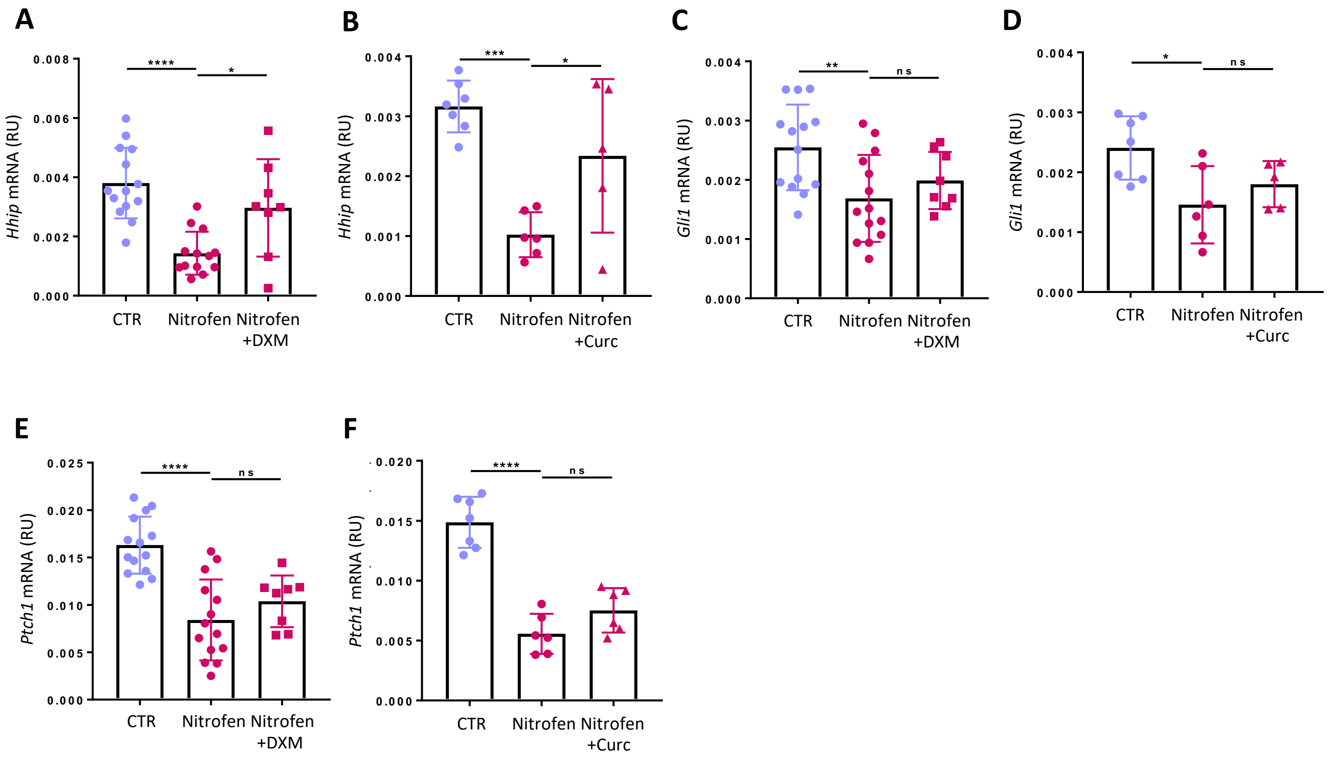
